# Cross-species prediction of essential genes in insects through machine learning and sequence-based attributes

**DOI:** 10.1101/2021.03.15.433440

**Authors:** Giovanni Marques de Castro, Zandora Hastenreiter, Thiago Augusto Silva Monteiro, Francisco Pereira Lobo

**Affiliations:** Universidade Federal de Minas Gerais, Belo Horizonte, Minas Gerais, Brazil

**Keywords:** *Drosophila melanogaster*, *Tribolium castaneum*, Essential gene prediction, Machine Learning, Homology-independent features, sequence-based attributes

## Abstract

Insects are organisms with a vast phenotypic diversity and key ecological roles. Several insect species also have medical, agricultural and veterinary importance as parasites and vectors of diseases. Therefore, strategies to identify potential essential genes in insects may reduce the resources needed to find molecular players in central processes of insect biology. Furthermore, the detection of essential genes that occur only in certain groups within insects, such as lineages containing insect pests and vectors, may provide a more rational approach to select essential genes for the development of insecticides with fewer off-target effects. However, most predictors of essential genes in multicellular eukaryotes using machine learning rely on expensive and laborious experimental data to be used as gene features, such as gene expression profiles or protein-protein interactions. This information is not available for the vast majority of insect species, which prevents this strategy to be effectively used to survey genomic data from non-model insect species for candidate essential genes. Here we present a general machine learning strategy to predict essential genes in insects using only sequence-based attributes (statistical and physicochemical data). We validate our strategy using genomic data for the two insect species where large-scale gene essentiality data is available: *Drosophila melanogaster* (fruit fly, Diptera) and *Tribolium castaneum* (red flour beetle, Coleoptera). We used publicly available databases plus a thorough literature review to obtain databases of essential and non-essential genes for *D. melanogaster* and *T. castaneum*, and proceeded by computing sequence-based attributes that were used to train statistical models (Random Forest and Gradient Boosting Trees) to predict essential genes for each species. Both models are capable of distinguishing essential from non-essential genes significantly better than zero-rule classifiers. Furthermore, models trained in one insect species are also capable of predicting essential genes in the other species significantly better than expected by chance. The Random Forest *D. melanogaster* model can also distinguish between essential and non-essential *T. castaneum* genes with no known homologs in the fly significantly better than a zero-rule model, demonstrating that it is possible to use our models to predict lineage-specific essential genes in a phylogenetically distant insect order. Here we report, to the best of our knowledge, the development and validation of the first general predictor of essential genes in insects using sequence-based attributes that can, in principle, be computed for any insect species where genomic information is available. The code and data used to predict essential genes in insects are freely available at https://github.com/g1o/GeneEssentiality/.

## Background

The set of genes where loss of function (LOF) mutations causes the death of an organism before reproduction or its lack of reproductive capabilities is defined as essential genes (Rancati et al. 2018). The discovery and characterization of essential genes has a broad range of scientific and technological applications. In cellular biology, the detection of the minimum set of genes for a species to thrive in specific environments defines the minimal genomes for specific organism/environment pairs, allowing both the definition of core molecular processes for life and the developmental of engineered organisms for a broad range of biotechnological applications (Hutchison et al. 2016). Essential genes are also interesting molecular targets for pharmacological intervention in pathogenic bacteria (Wang et al. 2014), insect pest management (Baum et al. 2007; Knorr et al. 2018) and cancer therapies (Chen et al. 2017). However, the experimental identification of essential genes on a large scale is a resource intensive task that may not be feasible for all organisms and genes.

At least partially driven by this limitation, several computational methods to predict gene essentiality in several species have already been developed, ranging from simple annotation transfer procedures to machine learning approaches (Acencio and Lemke 2009; Plaimas et al. 2010; Philips et al. 2017; Tian et al. 2018). The heterogeneous gene-centric features used in these strategies may be broadly classified in one out of two groups: 1) extrinsic properties that rely on external information, such as comparative genomics or experimental data; 2) intrinsic properties that can be computed from sequence data alone, such as statistical and physicochemical descriptors (Nigatu et al. 2017; Campos et al. 2020).

A major predictor of essential genes using extrinsic information is homology inference, consisting on the identification of genes in a species of interest that are putative homologs of known essential genes in other species and, consequentially, are likely to perform similar functions (Kumar et al. 2007; Holman et al. 2009). Another example of a phylogeny-based feature is the phyletic gene age, as older genes have a higher chance of being essential (Chen et al. 2012). Even though successful in several cases, homology-based, gene-level annotation transfer cannot be used to predict essential genes that do not have homologs already described as an essential gene in other organisms, such as lineage-restricted genes.

Other extrinsic attributes are the experimentally-based features obtained from the empirical characterization of genes, their transcripts and functional products, such as the topological properties obtained from protein-protein interaction networks (Acencio and Lemke 2009) or information about cellular localization and gene expression profiles (Cheng et al. 2014). Even though these properties have been widely used in machine-learning approaches to predict essential genes, they require expensive and laborious large-scale experimental data that are not readily available for non-model organisms.

Intrinsic features are defined as the sequence-based attributes that can be calculated from the gene sequence alone without relying on any external data. Representative examples are nucleotide, dinucleotide and codon compositions, as well as longer strings, information content, gene length, among others (Coutinho et al. 2015; Nigatu et al. 2017). For protein-coding genes, equivalent properties are also computable at the protein sequence level. Additionally, several protein-specific representation schemas based on the global frequency and the distribution of physicochemical and structural properties of amino acids are also available, providing numerical representations of protein sequences that are commonly used in structural and functional protein researches (Xiao et al. 2015). As sequence-based features may be, at least in principle, obtainable for every gene regardless of their homology relationships and in the absence of experimental data, they are interesting attributes to develop more general computational tools based on statistical learning to predict essential genes without relying on extrinsic properties.

Most statistical models to predict gene essentiality were built for prokaryotes and have used a combination of phylogenetic, experimental and sequence-derived features when training models (Dong et al. 2018). The bias towards prokaryotic organisms is mainly due to the feasibility of large-scale screening assays to search for essential genes in unicellular organisms when growing under distinct conditions, and also due to the availability of this information as structured databases, therefore providing a wealth of gene essentiality data for several bacterial species (Luo et al. 2014).

The discovery of essential genes is far more challenging in multicellular organisms, as testing different conditions *in vivo* may be not feasible due to cost and experimental issues. Complex multicellular eukaryotic lineages require several developmental gene expression programs to produce a viable organism, and those essential genes may be discovered only when evaluating these species at the organism level. Not surprisingly, the majority of computational methods to predict essential genes in eukaryotes use as training model the unicellular yeast *Saccharomyces cerevisiae*, which may be studied in a similar manner as a prokaryotic species (Dong et al. 2018). In metazoans, most statistical models to predict gene essentiality were trained using data produced from essays in cell lines from model organisms *Homo sapiens* and *Mus musculus* (Yang et al. 2014; Guo et al. 2017; Philips et al. 2017), with far less studies aiming at predicting essential genes in organisms associated with the multicellular lifestyle (Tian et al. 2018).

Insects are a highly diverse group of metazoans playing key ecological, medical and agricultural roles (Rust and Su 2012; Crespo-Perez et al. 2020). Strategies to identify potential essential genes in insects may reduce the time and resources spent to detect and characterize genes playing roles in core processes of insect biology. Furthermore, the prediction of essential genes that occur only in insects or in specific groups within insects, such as lineages containing insect pests and vectors, may provide a more rational procedure to select genes for molecular intervention with fewer off-target effects.

The fruit fly *Drosophila melanogaster*, arguably one of the most well-characterized multicellular model organisms, has approximately 52% of its protein-coding genes annotated with LOF phenotypes (Ewen-Campen et al. 2017). A useful resource to provide additional LOF information for *D. melanogaster* is the Flybase, which contains curated and structured data about known loci in the fruit fly, such as alleles, genotypes, and their associated phenotypes, allowing one to systematically survey this information for LOF alleles in specific genotypic configurations that results in phenotypes either incompatible with life or viable, and consequently obtain list of essential and non-essential genes, respectively, and including developmental essential genes (Larkin et al. 2021). Gene essentiality data in *D. melanogaster* is also available from genome-wide studies done in cell lines; however, this information probably does not capture the importance of developmental essential genes needed for multicellular species, as they rely mostly on large scale *in vitro* searches for essential genes at the cell level (Boutros et al. 2004; Viswanatha et al. 2018).

The second insect where genomic-scale gene essentiality information is available is the red flour beetle (*Tribolium castaneum*), an emerging model organism with an increasing amount of genomic and phenotypic information available. The iBeetle-base is a structured database containing curated information about gene silencing in *T. castaneum* using RNAi in different developmental stages, including lethality data (Donitz et al. 2015).

The insect orders Diptera and Coleoptera diverged approximately 300 million years ago, while insects evolved around 600 million years ago (Kumar et al. 2017). Consequently, even though these two orders share a considerable fraction of homologous genes, lineage-specific gene sets have evolved. It is reasonable to suppose that some of these lineage-specific genes code for biological functions needed for lineage-specific essential processes (Chen et al. 2010). Therefore, *D. melanogaster* and *T. castaneum* comprise an interesting scenario to develop and evaluate a general statistical workflow to predict essential genes in phylogenetically distant insects, including evaluation of performance while predicting lineage-specific essential genes.

The combined usage of extrinsic and intrinsic features has been demonstrated to improve prediction performance when searching for essential genes. Recently, two studies have demonstrated a considerable performance to predict essential genes in *D. melanogaster* by using both extrinsic and intrinsic information (Aromolaran et al. 2020; Campos et al. 2020). However, as previously exposed, extrinsic features have major drawbacks that prevents them to be used in general strategies to predict essential genes in insects. Due to the scarcity of experimental omics data for most insects, any approach to develop a general predictor of insect essential genes relying on experimental data will lack applicability in non-model organisms.

Here we describe a method to gather essential and non-essential protein-coding gene data for *D. melanogaster* and for *T. castaneum* from the Flybase and the iBeetle-base, respectively. We also obtained, for each protein-coding gene, a set of intrinsic gene- and protein-based features that were used to successfully train ML models for the prediction of essential genes in these two organisms. We demonstrated that ML models trained with *D. melanogaster* data can predict essential genes in *T. castaneum* and vice versa, suggesting our models could be used to predict essential genes in non-model insects that do not possess gene essentiality information. Finally, we demonstrated our *D. melanogaster* ML model can also detect essential genes in *T. castaneum* that have no homologs in the fly, demonstrating how our strategy of using intrinsic features for gene essentiality prediction allows one to survey gene sequences that would not be detectable using homology-based attributes. The source code and databases used in this study, as well as trained models that can be used to predict essential genes in other insect species, are freely available at https://github.com/g1o/GeneEssentiality/.

## Methods

### Dataset of essential and nonessential genes for *D. melanogaster* and *T. castaneum*

Our search for essential genes in Flybase was done using the query-builder tool. Two logical queries were built to unambiguously classify genes as either essential, non-essential or inconclusive (Figure 1A). The first query, called “Essential Gene Search” (EGS), aimed at finding essential genes, defined as those that have at least one LOF or hypomorphic allele in either homozygosity or heterozygosity associated with a lethal phenotype. The second query, called “Non-Essential Gene Search” (NEGS), aimed at finding non-essential genes, defined as those that do not have any LOF alleles with a lethal phenotype in homozygous individuals (Figure 1A).

**FIGURE 1:**
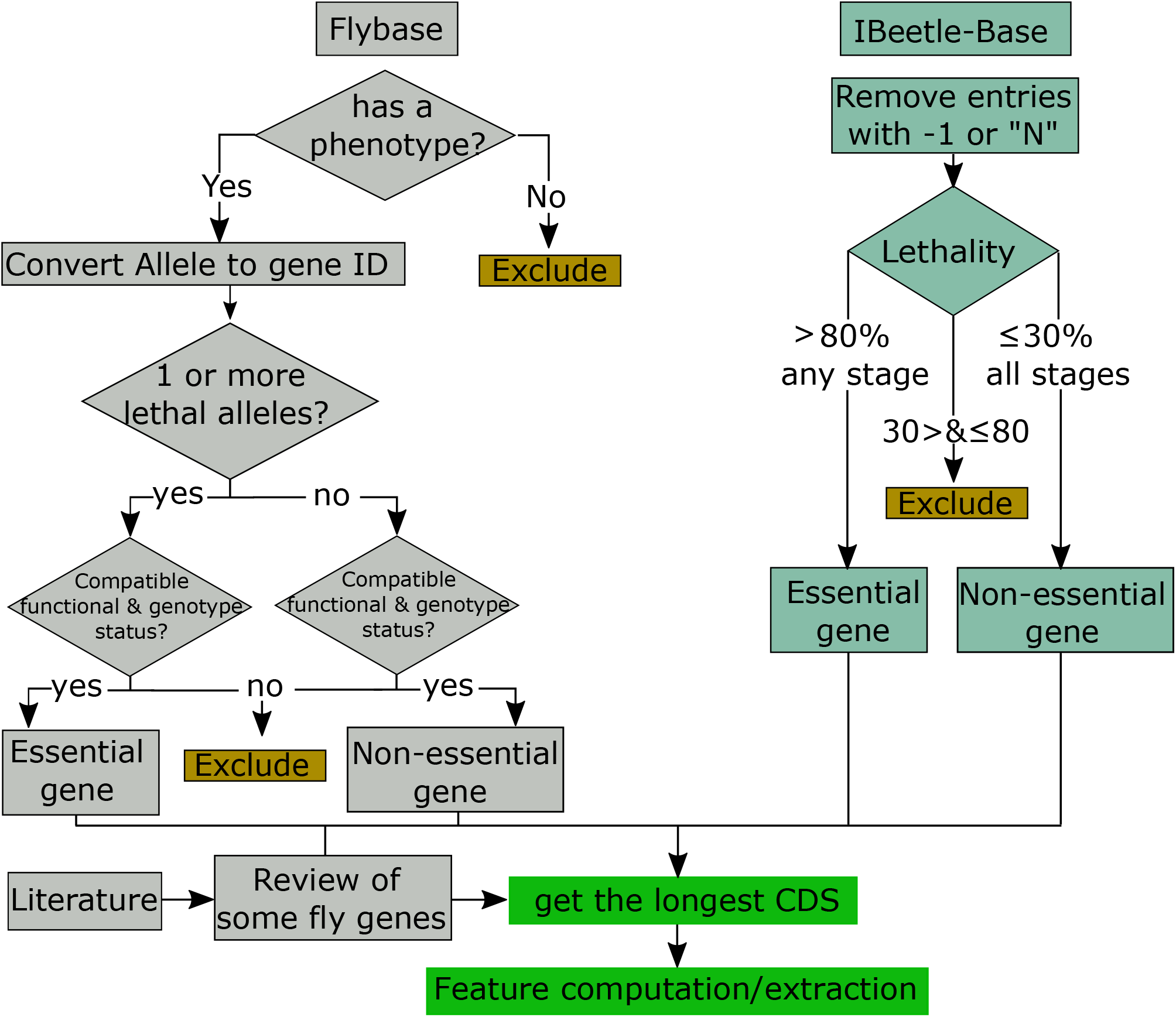
Flowchart for essentiality gene data acquisition. Essential and non-essential *D. melanogaster* genes selected during literature review were also included.

The EGS had the following parameters: The allele class had to be “loss of function allele” or “amorphic allele” or “amorphic allele - genetic evidence” or “amorphic allele - molecular evidence” or “hypomorphic allele” or “hypomorphic allele - genetic evidence” or “hypomorphic allele - molecular evidence” and the phenotypic class had to be “lethal”. Amorphic alleles are the ones where no gene function is reported, while hypomorphic alleles have their functions reduced in comparison to wild-type alleles. If a lower expression of an allele is enough to cause lethality, then we argue it is logical to classify this gene as essential.

As for the NEGS search, the parameters were the following: the allele class had to be “loss of function allele” or “amorphic allele” or “amorphic allele - genetic evidence” or “amorphic allele - molecular evidence”, but not “hypomorphic allele”, “hypomorphic allele - genetic evidence”, or “hypomorphic allele - molecular evidence”, and the phenotypic class had to be “not lethal”. We excluded hypomorphic alleles because they retain some biological activity by definition and, consequentially, do not provide information to objectively evaluate whether a LOF event for this locus is compatible with organism survival, which is the strict definition of non-essential genes.

In both searches, alleles containing no description of phenotypic class were excluded, as it is not possible to infer their essentiality status (Supplementary Table 1). No data from RNAi essays was selected for both searches, as these experiments do not provide allele class status. With the EGS and NEGS queries built and executed and proceeded by converting allele information to gene IDs.

To consider a gene as essential, we searched for alleles observed in either heterozygosity or homozygosity, as the LOF of a single allele is sufficient to detect dominant lethal alleles. Non-essential genes, however, can be safely classified as so only when a viable phenotype is observed in homozygous individuals, as recessive lethal alleles with LOF have been known for over a century [34, 35]. In *D. melanogaster*, the gene Duox is a classic example of a gene with recessive lethal alleles [36]. Therefore, we considered as non-essential genes only those where individuals with amorphic alleles in homozygosity are viable, as heterozygous viable individuals may either represent true non-essential genes or recessive lethal genes.

The genes found in both EGS and NEGS were considered as essential genes based on the premise that a gene having at least one LOF lethal allele is an essential gene in at least one condition (conditional essential genes). We evaluated our automatic query to recover essential and non-essential fly genes through an extensive literature review for genes already described as either essential or non-essential. This literature review also found essential and non-essential genes previously not categorized by our queries that were used to expand our dataset (Figure 1). The combined dataset of the Flybase search and the literature review will be referred from now on as DMEL dataset.

Previous works aiming at large-scale prediction of essential *D. melanogaster* genes relied on specialized databases of essential genes (OGEE and DEG) to gather gene essentiality status data, obtained mostly through large-scale *in vitro* searches through knockout and knockdown essays (Campos et al. 2019; Aromolaran et al. 2020). However, these datasets lack interesting properties found in FlyBase, such as the possibility of using functional, genotypic, allelic and phenotypic data to construct queries and select essential and non-essential genes, or the likely underrepresentation of developmental essential genes. Due to these reasons, we decided to use FlyBase as our single source of gene essentiality data (see Supplementary File 1). We compared our sets of essential and non-essential genes with the ones used in the work by Campos *et al*. (Campos et al. 2020), which also uses FlyBase as a source of essential and non-essential genes to train a predictor of essential genes for *D. melanogaster*.

The iBeetle database contains results of experiments from thousands of silenced genes in *T. castaneum* thought injection of dsRNA during both pupal and 5th/6th instar larval stage, including lethality data 11 days post injection (dpi) for pupal injection and 11 and 22 dpi for larval injection (Donitz et al. 2015). We queried the iBeetle database to obtain the information of gene lethality for the three developmental stages and computed the lethality distribution similarly as done by (Schmitt-Engel et al. 2015). Specifically, we required essential genes to have more than 80% lethality at 11 or 22 days after larval injection or 11 days after pupa injection of dsRNA. As for the non-essential genes, they were defined as those with at most 30% lethality at all time points, after both larval and pupal injection. This complete set of essential and non-essential genes from *T. castaneum* will be referred as TRIB dataset from now on. The iBeetle database also provides information about putative homologs in of *T. castaneum* genes in *D. melanogaster* as predicted from OrthoDB (Kriventseva et al. 2019). By selecting genes from TRIB dataset that had an empty OrthoDB field, we are selecting beetle genes that do not have a homolog in the fly genome (*T. Castaneum-specific* genes). From now on we refer to this dataset as noh-TRIB (no-homologs *Tribolium* dataset).

### Feature extraction

We obtained the longest coding region for each protein-coding gene to avoid potential biases caused by genes that have more than one transcript. The coding region was translated using the standard genetic code, and the features for each gene were extracted from both CDS and protein sequences using a combination of external software, R packages and *in-house* code (R_Development_Core_Team 2016) (Table 1 contains the summary of the features computed in this study, as well as the software used to calculate them). The source code for this analysis is available at https://github.com/g1o/GeneEssentiality/. Mutual Information (MI), Conditional Mutual Information (CMI) and Shannon Entropy (H) were calculated for DNA and protein sequences as done by Nigatu *et al*. (Nigatu et al. 2017). The other features present in Table 1 but not described below were extracted using functions from seqinR (Charif 2007), rDNAse (Zhu M 2016) and protR (Xiao et al. 2015) packages, and are described in the respective package documentations.

**TABLE 1:** Features extracted from nucleotide and amino acid sequences and used for classifier development.

| Feature | Sequence class | Number of features | Software/<br>package |
| --- | --- | --- | --- |
| **Mutual information** | Nucleotide | 17 | This work |
| **Conditional mutual information** | Nucleotide | 65 | This work |
| **Shannon entropy** | Nucleotide | 2 | This work |
| **Dinucleotide-based Auto-cross Covariance** | Nucleotide | 2888 | rDNase |
| **Trinucleotide-based Auto-cross Covariance** | Nucleotide | 2100 | rDNase |
| **Pseudo Dinucleotide Composition** | Nucleotide | 19 | rDNase |
| **Gibbs entropy and enthalpy** | Nucleotide | 2 | VarGibbs |
| **Physicochemical classes & Theoretical isoelectric point** | Amino acid | 10 | Seqinr |
| **Mutual information** | Amino acid | 41 | This work |
| **Conditional mutual information** | Amino acid | 8001 | This work |
| **Shannon entropy** | Amino acid | 2 | This work |
| **Protein length** | Amino acid | 1 | This work |
| **Conjoint triad** | Amino acid | 343 | Protr |
| **MoreauBroto** | Amino acid | 240 | Protr |
| **Moran** | Amino acid | 240 | Protr |
| **Geary** | Amino acid | 240 | Protr |
| **CTD Descriptors - Composition (CTDC)** | Amino acid | 21 | Protr |
| **CTD Descriptors - Distribution (CTDD)** | Amino acid | 105 | Protr |
| **CTD Descriptors - Transition (CTDT)** | Amino acid | 21 | Protr |
| **Pseudo amino acid composition** | Amino acid | 52 | Protr |
| **APSEAA** | Amino acid | 120 | Protr |

#### Gibbs Entropy

Each nucleotide sequence was used as an input to VarGibbs (Weber 2015), which predicted the Gibbs entropy and enthalpy for DNA sequences using the model D-CMB and produces two features as output.

#### Pseudo-amino acid composition

The use of individual amino acid frequency fails to consider the possible interactions that may exist between them by discarding information about amino acid order in protein sequences. The pseudo-amino composition was developed to represent this information as numerical features, and later on used for tasks such as the prediction of cellular location and membrane proteins (Chou 2001). There are two main parameters needed to calculate pseudo-amino acid composition: lambda and omega. The lambda is the maximum distance between the amino acid residues in protein sequences to be considered, and is used to calculate the sequence order correlation. Lambda values determine the number of features to be generated, and it needs to be at least of the length of the protein sequence, as it generates 20 + lambda features; the first 20 features reflect the effect of amino acid composition, while the additional ones describe the sequence order effect. We used a lambda value of 50. The omega is the weight factor for the sequence order effect, and for this we used the default value (0.05). The EXTRACT_PAAC function from protR was used to calculate this feature. The non-coding genes and proteins that did not have the required length of 50 amino acids to calculate the pseudo-amino acid features were excluded from downstream analyses.

#### Protein length

The length of the protein is a simple attribute already used by other ML models to predict essential genes, which tend to be have a greater length than non-essential ones (Grishkevich and Yanai 2014). We used built-in functions from the R language to compute sequence length.

#### Conjoint Triad

Initially used to predict protein–protein interactions, this feature classifies the 20 amino acids into 7 classes according to their physicochemical properties, proceeding by counting the frequencies of each three sequential amino acids residues according to their classes, resulting in a total of 343 features (Shen et al. 2007). We used the extractCTriad function from protR to produce conjoint triad features.

### Model training and cross-validation

We computed the sequence-based features described in Table 1 and used them to train and evaluate statistical models capable of distinguishing between essential and non-essential genes. We started by applying a zero-variance filter to remove constant features and used the remaining features as input to train Extreme Gradient Boosting Trees (XGBT) and Random Forest (RF) models using the methods ‘xgbTree’ (Chen T. 2016) and ‘ranger’ (Wright 2017) from the caret package (Kuhn 2008) (Supplementary Figure 1). The parameters for the RF models were 1000 trees and tuning by selecting the *mtry* between the squared number of features and two times itself. As for the XGBT models, the parameters held constant were *eta* = 0.1, *gamma* = 1, *colsample_bytree* = 1, *min_child_weight* = 1, *subsample* = 1 and, for parameter tunning, *nrounds* was 100, 200 and 500 and *max_depth* was 4 and 10.

The ten-fold cross-validation used when training the models was repeated 3 times, using the maximum AUC of the ROC (AUC-ROC) as the performance metric to be maximized. We compared the performance of distinct models using the DeLong’s test as implemented in the roc.test function from the pROC package (Sun 2014). Specifically, we were interested in evaluating 1) the performance of predictors of essential genes in *D. melanogaster* and *T. castaneum* using only intrinsic attributes and 2) the influence of model type on classifier performance (random forest versus extreme gradient boosting). To evaluate the performance of insect-specific models against a baseline, we used a zero-rule (ZR) classifier that classifies every protein as the majority one. Even though we do not have a highly unbalanced training dataset, we also report precision-recall AUCs (PR-AUCs) for both trained and ZR models to evaluate classifier performance in terms of false positive and false negative rates.

### Model generalization through transfer learning: interspecies test and *T. castaneum-restricted* genes

To evaluate if the models trained using *D. melanogaster* or *T. castaneum* data are general classifiers that can predict essential genes in other insect species, we used each model to predict essential genes in the other insect species (Supplementary Figure 1). To check if the *D. melanogaster* model can predict essentiality in *T. castaneum*-restricted genes (genes that have no homologs in *D. melanogaster*), we used the fly models to predict essential genes in the noh-TRIB dataset. Again, the ROC-AUCs were tested for a significant difference against each other and the result of the ZR model to evaluate whether the AUCs obtained in these tests are significantly better than null models.

## Results/Discussion

### Database of essential genes and gene-centric attributes for *D. melanogaster* and *T. castaneum*

We built two queries to survey the Flybase database and select sets of alleles where specific combinations of genotypic and functional information about alleles in a locus unambiguously resulted in either a lethal (essential genes) or viable (non-essential genes) phenotype, after excluding inconclusive cases (Figure 1; see also “Methods” section). In our search, we observed 1267 alleles with no described phenotypic class that were excluded from downstream analysis, as they could not be automatically evaluated regarding their essentiality phenotype (Supplementary Table 1).

After mapping and filtering the alleles from the EGS and NEGS to gene IDs, we obtained 1393 essential and 899 non-essential genes. A literature review found 38 and 56 essential and non-essential genes, respectively, also present in our Flybase query results. From these, 32 (84%) and 53 (95%) genes were correctly classified as essential and non-essential genes by our query, respectively, while 6 (16%) essential and 3 (5%) non-essential genes were false positives and negatives from our query, respectively (precision of 0.84, recall of 0.91 and F-measure of 0.87). Therefore, we concluded our query is capable of automatically selecting sets of essential and non-essential genes.

During the literature review, we also found 13 essential and 37 non-essential genes not present in our Flybase query that were added to our dataset, resulting in 145 reviewed genes (52 essential and 93 non-essential genes, Supplementary Table 2). After merging Flybase and our literature review data, our *D. melanogaster* dataset had 1406 essential and 936 non-essential protein-coding genes. We proceed by selecting only protein-coding genes, which resulted in our final dataset of containing 1333 and 753 essential and non-essential genes, respectively (DMEL dataset).

The DMEL dataset produced in our analysis shares similarities but also has considerable differences compared with the dataset used by Campos *et al.* (2020) (Supplementary Figure 2). The majority of our essential genes (~ 91%, 1213 out of 1333) are classified by Campos *et al.* (2020) as either essential (109) or conditional essential (1104). A similar pattern was also found for our non-essential genes, which were mostly classified as non-essential (486) and, in fewer observations, as conditional essential genes (187). We observed 43 and 10 genes classified as essential and non-essential in our analysis, respectively, that were classified as non-essential and essential by Campos *et al.* (2020). These authors also report a considerable number of essential (295), conditional essential (2991) and non-essential (6355) genes not selected in our searches in FlyBase, while our search found 47 and 40 exclusive essential and non-essential genes, respectively.

We suggest these differences to be caused by several factors. One difference that may account for the larger number of genes exclusively available in Campos and collaborators dataset is the fact that we use both phenotype and allele class information, together with genotype status per locus, to objectively classify genes as either essential or non-essential. A considerable fraction of loci is inconclusive after taking into account these known confounding biological factors, and this data is removed from downstream analyses. As an example, we mention the high fraction of alleles missing allele class information in *D. melanogaster* genes (Supplementary Figure 1, Supplementary Table 1).

Campos *et al.* (2020) used all the genes where alleles with phenotypic status identified as either “lethal” or “viable” are available to compute an *ad-hoc* essentiality score (ES) for each of these genes, defined as the total number of alleles linked to essential/lethal terms squared divided by the total number of experiments linked to essential/lethal plus non-essential/viable terms squared. The ES score was then used to classify every gene in one out of three categories: essential (ES > 0.9), conditional essential (0/9 > ES > 0.1) or non-essential (ES < 0.1). Therefore, the very concept of essentiality, and the strategies used to define it, are substantially distinct in our studies, which are likely to produce distinct gene sets from the beginning.

Our definition of essential genes (genes with LOF or hypomorphic alleles causing lethal phenotypes in either heterozygosity/homozygosity regardless of environmental conditions) tries to translate the broadest logical definition of essential genes while taking into account genotypic, phenotypic and functional data, and certainly includes conditional essential genes. This fact is coherent with the distribution of our essential genes according to the ES by Campos *et al.* (2020), if we assume the majority of our essential genes are classified as conditional essential by them.

The definition of non-essential genes in our work – loci where LOF alleles in homozygosity result in viable individuals – also takes into account genotypic, phenotypic and functional data to exclude data that may prevent one to unambiguously classify a gene as non-essential, such as the occurrence of recessive lethal and hypomorphic alleles. We argue these differences may be at least partially responsible for the larger number of genes exclusively classified as non-essential by Campos *et al.* (2020) compared to our data. Their score-based strategy would accept as non-essential, for instance, the loci where only a small fraction of alleles causes lethality, which may even include essential genes with lower penetrance (Schmitt-Engel et al. 2015).

As for *T. castaneum*, the iBeetle-base contains information about lethality for 4084 genes evaluated at 11 dpi for pupal (P11) and larval (L11) stage injection and at 22 dpi for the larval (L22) stage, while 4174 genes having missing data in the lethality field and were excluded from downstream analysis (Figure 1). When evaluating the relative frequency of genes as a function of lethality in the three developmental stages, we found it to be biased towards either genes with high or low lethality values (Figure 2). After a visual inspection of our histograms, and also as previously used by (Schmitt-Engel et al. 2015), we defined essential genes as the ones with a lethality value greater than 80% in at least one developmental stage (Figure 2, purple dots) and as non-essential genes the ones with a lethality value smaller than or equal to 30% in all three developmental stages (Figure 2, yellow dots).

**FIGURE 2:**
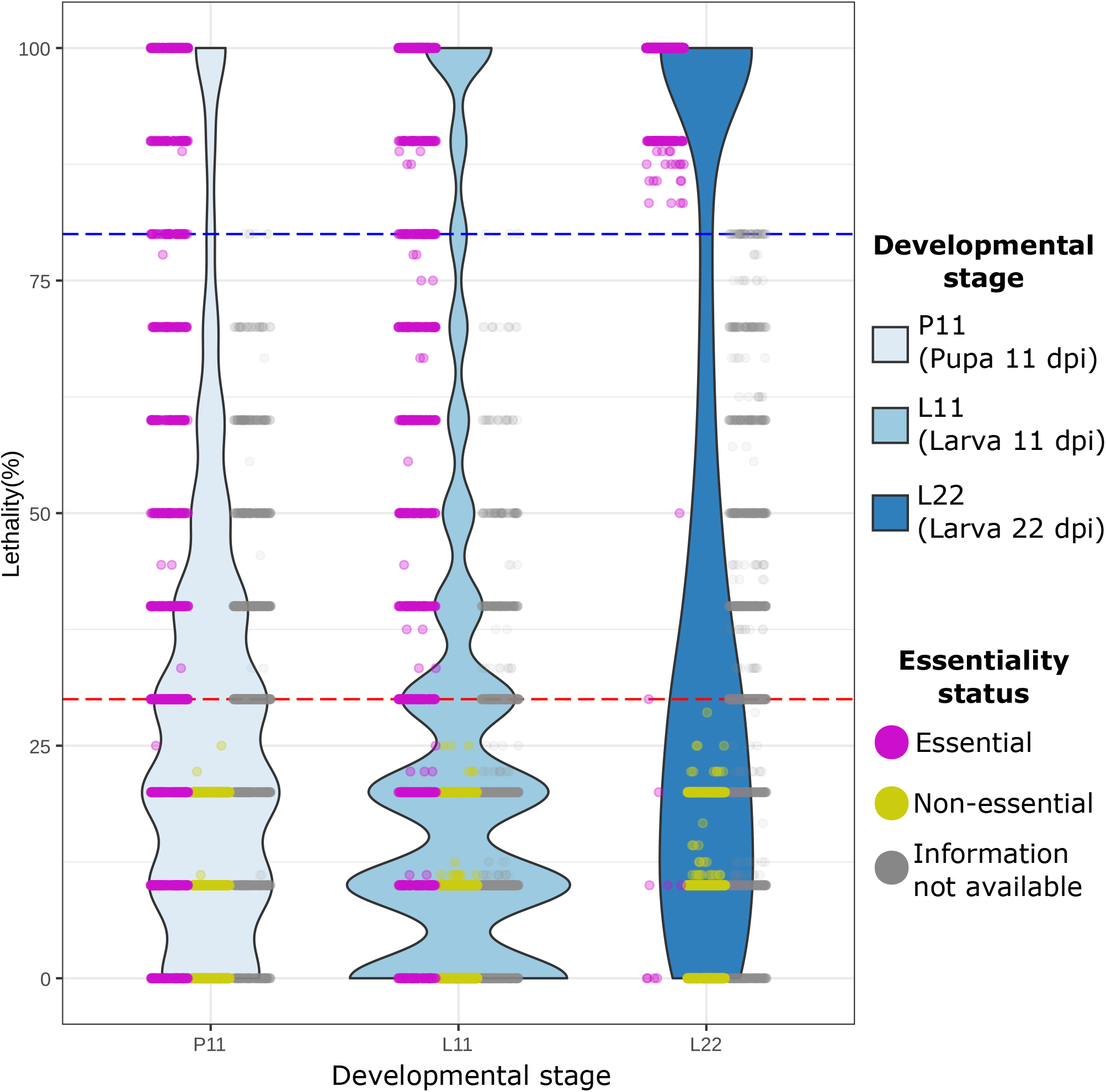
Distribution of *T. castaneum* genes using the lethality data from iBeetle. P11: Pupa 11 days post injection; L11 and L22: Larva 11- and 22-days post injection, respectively. The dots show genes classified as essential (purple), non-essential (yellow) or those that could not be classified (grey).

We observe a wide distribution of lethality values for essential genes in the larval and pupal developmental stages 11 dpi (L11 and P11) with a slight increase in lethality values around 100% (Figure 2, purple dots). This contrasts with the much higher mortality of essential genes in the only developmental stage where lethality data 22 dpi is available (larval 22 dpi, L22). This distribution suggests that there may be relatively few housekeeping essential genes with 100% mortality in distinct developmental stages before 11 days. On the other hand, the majority of essential genes required at least 12 days to be detected as so in the larval stage, a phenomenon likely caused by low-penetrance lethal phenotypes (Schmitt-Engel et al. 2015). After excluding two genes with conflicting lethality information, we selected 1073 and 1077 essential genes and non-essential genes for *T. castaneum*, respectively. Our search for *T. castaneum*-restricted genes found respectively 35 and 104 essential and non-essential genes with no homologs in *D. melanogaster* (Figure 1; see also “Methods” section).

### Prediction of essential genes in *D. melanogaster* and *T. castaneum* using species-specific classifiers

We used the DMEL and TRIB datasets and the XGBT and RF models to initially train four classifiers that were evaluated to determine whether it is possible to predict essential genes in distinct insect species by using exclusively intrinsic attributes, and also to evaluate relative model performance. Both models could distinguish between essential and non-essential genes in the two insect species with performances significantly better than the ZR models (Figure 3, p-value < 1e-10 for all model comparisons). The ZR model has a fixed ROC-AUC = 0.5, as it classifies every gene with the same probability, but its precision recall curve (PRC) is dependent on the relative frequency of true positives, therefore producing different results for each dataset.

**FIGURE 3:**
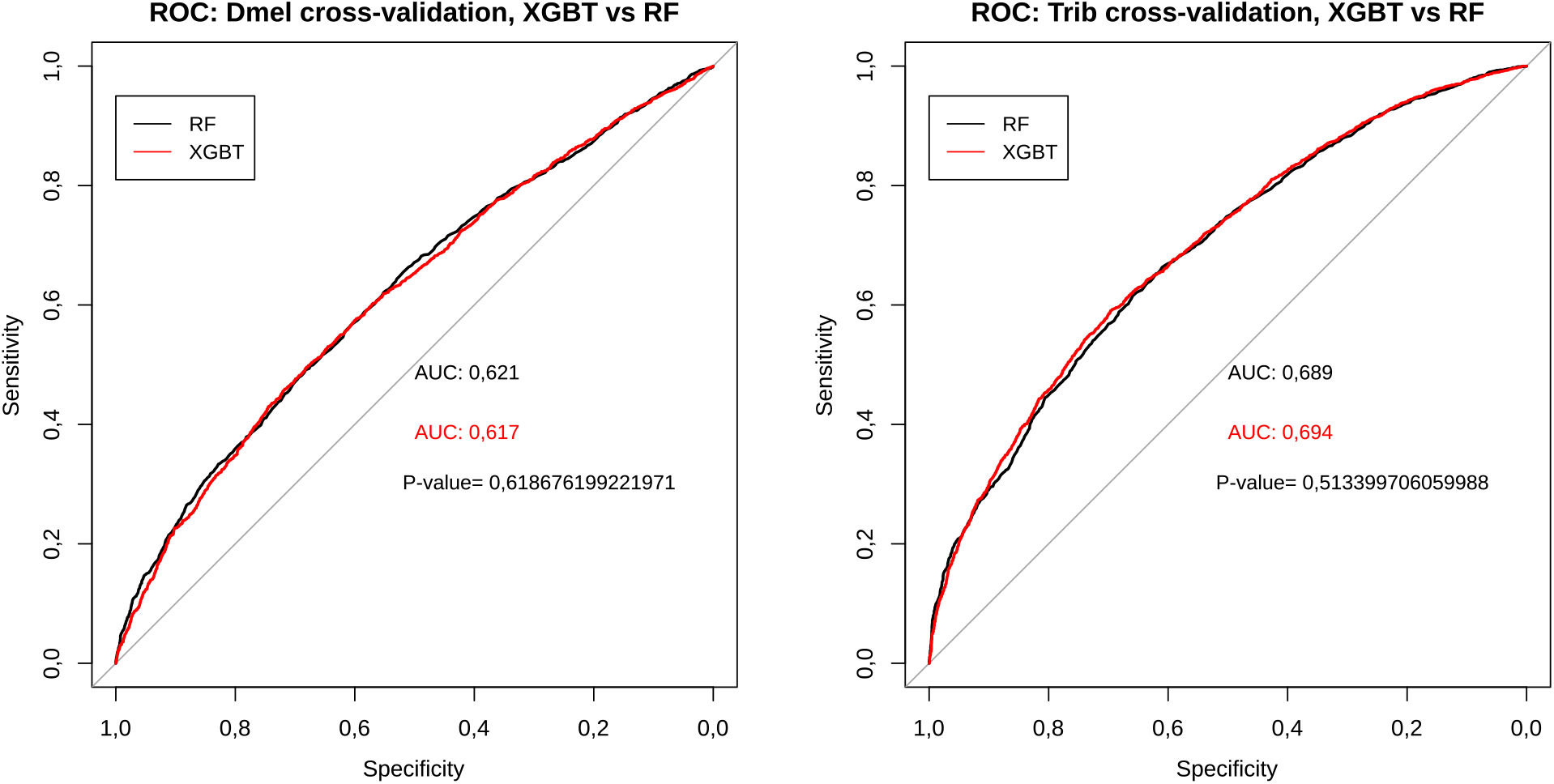
Cross-validation data from the XGBT and RF models. Black curves: Random Forest models, red curves: XGBT models. A) *D. melanogaster* model. B) *T. castaneum* model.

Regarding relative model performance, we found the cross-validation of *D. melanogaster* models to have ROC-AUC values of 0.618 and 0.621 for the RF and the XGBT models, respectively while the *T. castaneum* cross-validation models have ROC-AUCs of 0.689 and 0.694 for the RF and XGBT models, respectively, indicating that both models have equivalent performance in both insect species, with no significant difference between them (p-value > 0.5 for all tests, Figure 3).

### Relative feature importance

We extracted the 20 most frequent features from the *D. melanogaster* and *T. castaneum* RF models to qualitatively evaluate the properties of (Supplementary Tables 3 and 4). We found both DNA and protein attributes to be present among the most important features for the two insect species, with DNA features corresponding to the majority of them (95% and 75% in Dmel and Trib models, respectively). The most important feature for the Dmel RF model was the B.DNA.twist.Roll.lag.1, a feature from the Dinucleotide-based Auto-cross Covariance.

As for the Trib RF model, the most important feature was the secondarystruct.Group2, extracted from the composition, transition and distribution (CTD) descriptors (Dubchak I. 1995). In brief, these features evaluate the global composition of amino acid catalogued to distinct attribute classes expected to reflect physicochemical and structural properties in sequences (e.g. the frequency of amino acids predicted to be in secondary structures of the class “strand”, as is the case for secondarystruct.Group2), the change of these properties across the sequence (e.g. the frequency of transitions from secondary structure “strand” to “helix” and vice-versa), and the global distribution of these properties along the sequences (e.g. the distribution of amino acids with attribute “helix” in the sequence).

Interestingly, even though the most relevant features are DNA-based properties that are not shared across the fly and beetle models, the single common feature for both models is a protein-derived one (prop1.Tr2332), comprising another CTD descriptor that represents the frequency of transitions from neutral to hydrophobic amino acids across protein sequences and vice-versa.

### Model generalization through transfer learning: interspecies test and *T. castaneum-restricted* genes

We further evaluated our general predictor of essential genes in insects by simulating a scenario of predicting essential genes in an insect species of interest using a pre-trained classifier using data from a phylogenetically distant insect. For that purpose, we used a leave-one-organism-out strategy, where training was done in one insect species and model testing was done int the other species not previously used to train the classifier.

Both XGBT and RF models, when trained using *D. melanogaster* data, are capable of predicting essential genes in *T. castaneum* significatively better than ZR model, with an ROC-AUC of 0.625 for the RF model (p-value = 1.85e-25) and ROC-AUC of 0.593 for the XGBT model (p-value = 1.85e-25) (Figure 4A), and with precision-recall AUC (PRC-AUC) values of 0.606 and 0.581 for the RF model and XGBT models, respectively, while the ZR model had a PRC-AUC of 0.499 (Figure 5 A). A similar scenario was observed for both model classes when training was done using *T. castaneum* data and model evaluating was done using *D. melanogaster* data compared to a ZR model. We observed ROC-AUC values of 0.606 for the RF model (p-value = 1.72e-16) and of 0.597 for the XGBT model (p-value = 5.59 e-14, Figure 4B), and PRC-AUC values of 0.713 and 0.709 for the RF and XGBT models, respectively, while the ZR model had a PRC-AUC of 0.639 (Figure 5B).

**FIGURE 4:**
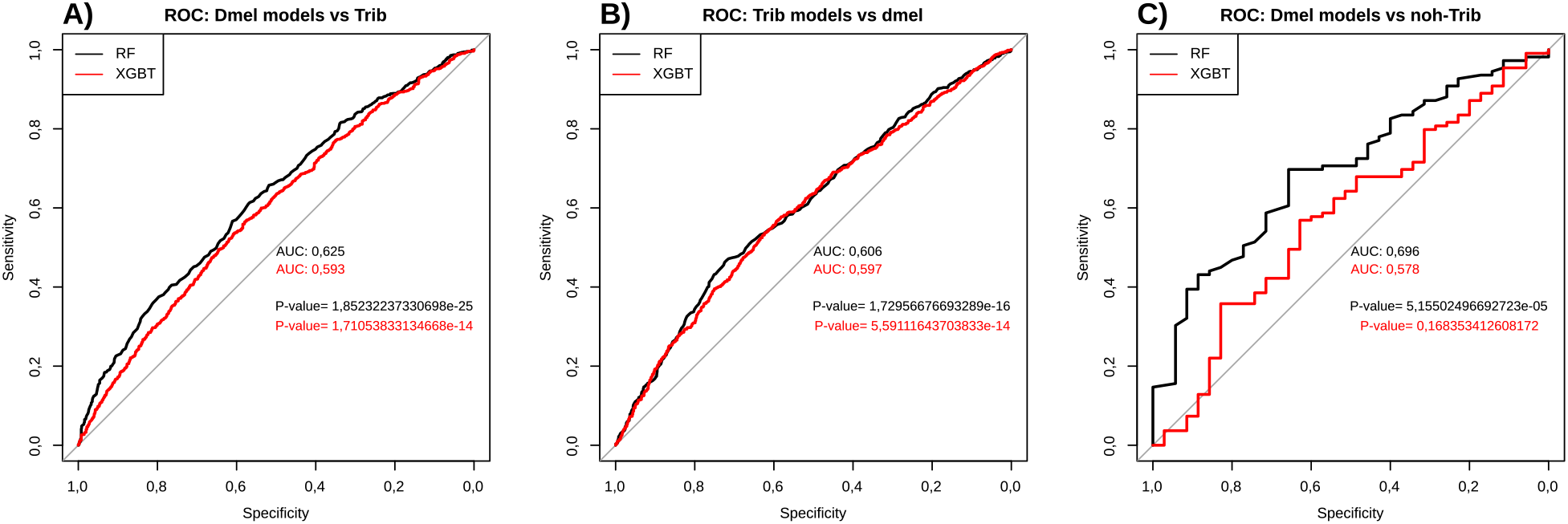
Cross-validation data for model evaluation in distinct scenarios. The gray line is the ROC for the zero-rule model (ZR) where all genes are classified as essential with the same probability of 1. By definition, the ROC-AUC of the ZR model is 0.5. Black curves: Random Forest models, red curves: XGBT models. A) Prediction of *T. castaneum* genes using Dmel models. B) Prediction of the *D. melanogaster* genes using the Trib model. C) Prediction of *T. castaneum* genes that have no homologues in *D. melanogaster* using the Dmel model.

**FIGURE 5:**
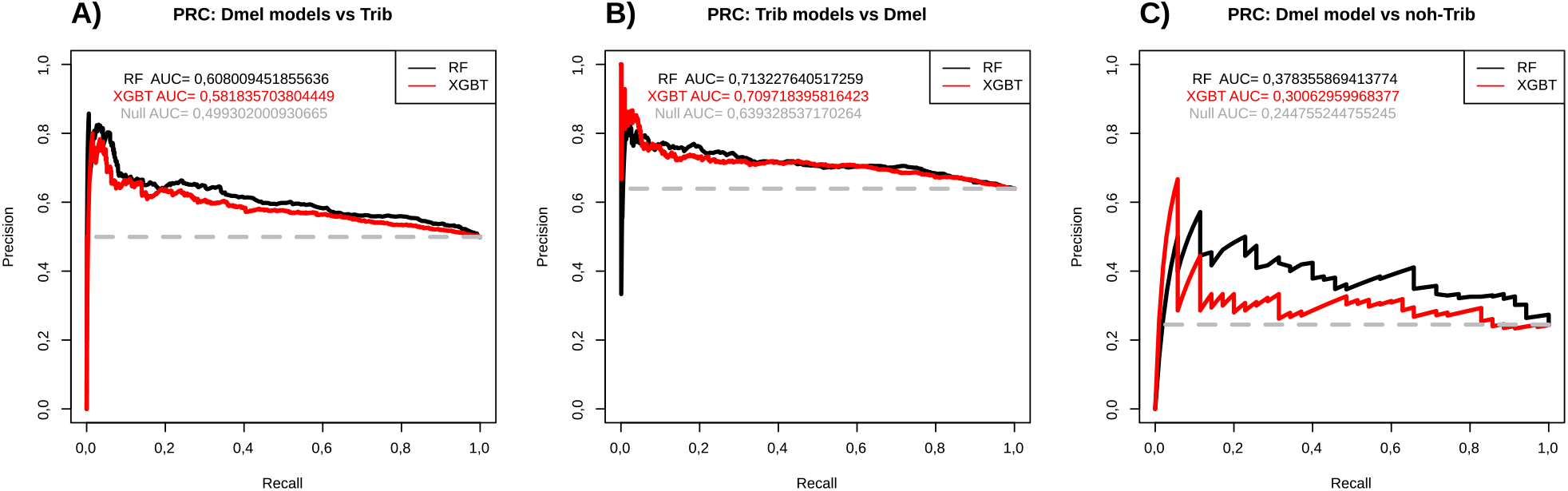
Precision recall curves (PRC) for the test sets. The dashed red line is the PRC for the ZR model, in which all genes are classified as essential with the same probability of 1. AUC-PRC of the ZR model is proportional to the frequency of true positives, as its value depends on the test set. A) Prediction of *T. castaneum* genes that have no homologues in *D. melanogaster* using the Dmel model. B) Prediction of all *T. castaneum* genes using the Dmel model. C) Prediction of the *D. melanogaster* genes using the Trib model.

At this point we conclude that both model classes (RF and XGBT), when trained in one insect species, can be used to predict essential genes in a phylogenetically distant insect species from another order, with performance significantly better than ZR models. Furthermore, even though the essential gene predictors selected distinct features when training using data from distinct insect species, they are still capable of achieving a cross-species classification performance significantly better than expected by chance.

As a final experiment we evaluated whether our *D. melanogaster* models can predict essential *T. castaneum* genes that have no known homolog in the fly (noh-TRIB dataset, comprising 35 essential and 109 non-essential genes). The *D. melanogaster* RF model had an ROC-AUC=0.696, being significantly better than a ZR model (p-value = 5.15e-05), while the XBGT model had an ROC-AUC of 0.578, which is not significantly better than a ZR model using a p-value threshold of p < 0.05 (p=0.16, Figure 4C), while PRC-AUC values were 0.378 and 0.300 for the for the RF and XGBT models, respectively, and 0.244 for the ZR model (Figure 5C). Together, these results suggest it is possible to use our RF fly model to predict taxon-restricted essential genes in the beetle, which may be a useful resource to detect essential genes exclusively observed in some insect lineages.

## Conclusion

Insects are a group of organisms with an immense taxonomic diversity and over a million estimated species. This taxon is also highly diverse in terms of its ecological roles and importance to human societies, with several species observed as pollinators of crops and wild species, vectors of human and animal diseases, parasites, model organisms for basic and applied research, important components of nutrient cycles and food chains, and producers of economically relevant substances (Stork 2018). Therefore, it is highly desirable to be able to identify essential genes for specific groups of insects that can be used both to understand the molecular *modus operandi* of this taxon and also to develop lineage-restricted insecticides for pest control with potentially smaller ecological footprint.

Recently, two groups reported the successful development of predictors of essential genes for *D. melanogaster* (Aromolaran et al. 2020; Campos et al. 2020). Even though both methods are highly successful to predict essential genes for the fly, their usage as a general approach to predict essential genes in other insect species is highly limited. Both strategies used extrinsic gene-level properties, such as protein domain prediction, gene expression profiles and protein-protein interaction networks, to cite a few, together with intrinsic features, when developing their predictors. Although this information is readily available for the model organism *D. melanogaster*, this is not the case for the vast majority of insect species in a foreseeable near future, including *T. castaneum.* Not surprisingly, and in contrast to our work, both studies lack the formal demonstration that their models trained in *D. melanogaster* and using extrinsic features can be used to predict essential genes in other insect species.

Our work reports, to the best of our knowledge, the development and validation of the first general predictor of essential genes demonstrated to work in distinct insect taxa. We also report the compilation of sets of essential and non-essential genes for the two insect model organisms where this information is available – the fruit fly *D. melanogaster* and the red flour beetle *T. castaneum.* For *D. melanogaster*, FlyBase provides a curated dataset of genotypic and phenotypic information as produced by the entire fruit fly genetics community using a controlled ontology, therefore allowing one to construct logical queries to select gene sets containing only essential and non-essential genes. Most of these genes were surveyed for essentiality through small scale *in vivo* experiments performed by distinct research groups using individual knockouts and gene silencing experiments, therefore decreasing the chance of systematic biases introduced by error-prone high-throughput methods to search for essential genes such as transposon mutagenesis data (de Jong et al. 2014). As for *T. castaneum*, all essentiality data was obtained from large-scale injection of dsRNA targeting specific transcripts in experiments comprising distinct life stages (larval and pupal).

Our datasets of essential and non-essential genes for the two insect species were obtained using distinct experimental and search strategies, which decreases the possibility of systematic biases introduced by experimental variables (e.g., detection of cellular-level essential genes only being caused by using exclusively *in vitro* experiments to search for essential genes). Specifically, the experimental procedures used to generate essentiality data for both organisms comprise *in vivo* assays in several developmental stages, therefore allowing us to train models eventually capable of detecting developmental essential genes in addition to the essential genes corresponding to cellular-level processes. The large-scale gene essentiality information available for species *D. melanogaster* (Diptera) and *T. castaneum* (Coleoptera) and compiled in this work comprises a useful resource to develop and validate other predictors of insect essential genes, including lineage-specific candidates.

We proceed by demonstrating that models trained exclusively using sequence-based attribute data from one insect species and evaluated using gene sets from the same species have performance significantly better than expected by chance. Additionally, models trained using data from one insect species are capable of predicting essential genes in another insect species never used to train the model. This demonstrates the feasibility of essential gene prediction for insect species lacking this information by using models trained in phylogenetically distant insect species where phenotypic information about gene essentiality is available. We extended our analysis be demonstrating that the RF model trained in the fly successfully detect beetle-specific essential genes, therefore evidencing that even younger, lineage-restricted essential genes share enough common sequence-based properties with essential genes from a phylogenetically distant insect species to allow cross-species prediction of lineage-specific essential genes.

One can avoid using extrinsic features and still attain success when developing machine-learning strategies to predict essential genes in unicellular species (Guo et al. 2017; Liu et al. 2017; Nigatu et al. 2017; Campos et al. 2019). However, these studies have not shown how their predictive models behave when evaluating complex eukaryotic organisms or taxon-restricted genes. We found that Dmel and Trib RF models share a single feature across the most relevant ones, demonstrating that distinct combinations of features are being selected during model training. Nevertheless, these distinct feature sets are still capable of predicting essential genes in the other insect species. These facts, together with the performance values observed for our models, suggest we may be able to further improve classifier performance through feature engineering and by exploring other machine learning strategies. Our datasets and models can be used both as a tool to predict essential genes in insects and as benchmarks for further developments, and are freely available at https://github.com/g1o/GeneEssentiality/.

## Acknowledgment

This study was financed in part by the Coordenação de Aperfeiçoamento de Pessoal de Nível Superior - Brasil (CAPES). Thanks for the postgraduate programs of Genetics (PPG-Genética) and Bioinformatics (PPG-Bioinfo) of the Universidade Federal de Minas Gerais (UFMG). We would like to acknowledge the Sagarana HPC cluster/CEPAD/ICB/UFMG for the computational infrastructure, and Dr. Felipe Campelo França Pinto (Aston University/UK) for the fruitful discussions on machine learning.

