## Supplementary File 1, Supplementary Figures 1-2, Supplementary Tables 1-4 for "Cross-species prediction of essential genes in insects through machine learning and sequence-based attributes"

### Supplementary material

##### *D. melanogaster* data in OGEE, DEG and Flybase

The quality of the data used in machine learning is of utmost importance, as missing and mislabeled data for the training set may introduce systematic biases in models that are likely to decrease model performance [1]. In order to develop a predictor of essential genes for insects, we initially searched for large-scale experiments in *D. melanogaster* or databases describing both essential and non-essential genes.

We found this information available in two specialized databases containing lists of essential genes of *D. melanogaster* directly available – OGEE [2] and DEG [3]. OGEE contains two datasets for *D. melanogaster*: 1) 13781 genes surveyed for essentiality at the cellular level through *in vitro* RNAi knockdown [4], 2) 437 genes silenced with transgenic RNAi knockdown in flies at different developmental stages [5]. In DEG, we found one dataset containing 339 essential genes which were disrupted by P-element insertions [6]. The Flybase database uses controlled descriptions to describe allelic-centered data, such as mutation types, their functional consequences, and the associated phenotypes for distinct genotypes [7]. This structured information allows the selection of essential and non-essential genes through specific queries to unambiguously select such sets (See "Methods" for our query strategy).

We argue the OGEE and DEG datasets suffer from several drawbacks that undermines their effectiveness to produce trustable sets of essential and non-essential genes. First, none of these datasets report enough data to allow us to unambiguously select sets of essential and non-essential genes, as they lack the detailed allele-centered data needed to select such gene sets. Furthermore, the OGEE dataset 1 probably does not report developmental essential genes, falsely classifying them as non-essential genes, while OGEE dataset 2 was latter shown to contain results that might not be reproducible [8].

A deeper analysis of the alleles represented in the DEG dataset and in FlyBase found the majority of them to lack allele class information (86,7%, 1183 out of 1364). As for the 181 alleles where allele class information is available, we found 90 hypomorphic, 2 with gain of function and 1 hypermorphic allele, which prevents us to use it as a trusted source of LOF and hypomorphic alleles, which are fundamental to define essential and non-essential genes.

We also investigated if the dataset of essential genes we obtained with our Flybase search agrees with the largest available from OGEE, containing 267 essential genes from a total of 13781 tested genes [4]. This highly unbalanced dataset, containing 98,1% of the genes classified as non-essential, is likely to contain only housekeeping essential genes detected through *in vivo* high throughput screenings. In comparison, our search in Flybase resulted in 1393 essential genes and 899 non-essential genes. The intersection between non-essential genes from OGEE and essential genes from our search revealed that over 1000 essential genes from Flybase would be considered ad non-essential by this OGEE dataset.

After considering all aforementioned facts, we decided Flybase to be the best source of the genetic, functional, genotypic and phenotypic data, including data from several experimental sources and developmental stages, to unambiguously define sets of essential and non-essential genes. Therefore, we decided to proceed using this database as our source of gene essentiality data for *D. melanogaster*.

### Supplementary Legends

**SUPPLEMENTARY FIGURE 1: Flowchart for the model training and testing.**

**SUPPLEMENTARY FIGURE 2: Protein-coding essential (E), non-essential (NE), and conditionally essential (CE) genes from Campos et. al. (2020) compared with those found by us.** We used genes with an annotated LOF allele class and phenotype in addition to those we manually reviewed, while Campos group used genes based on alleles phenotypes.

**SUPPLEMENTARY TABLE 1: Most frequent terms of the phenotypic class from the NEGS search.** Alleles with an empty field were excluded.

**SUPPLEMENTARY TABLE 2: Manual curation of LOF alleles for *D. melanogaster*.** Each gene had at least one published research using a genetic knockout or null mutant that supports a lethal (essential) or viable (non-essential) phenotype. For the non-essential classification, LOF alleles had to be in a homozygous state.

**Supplementary Figure 1**

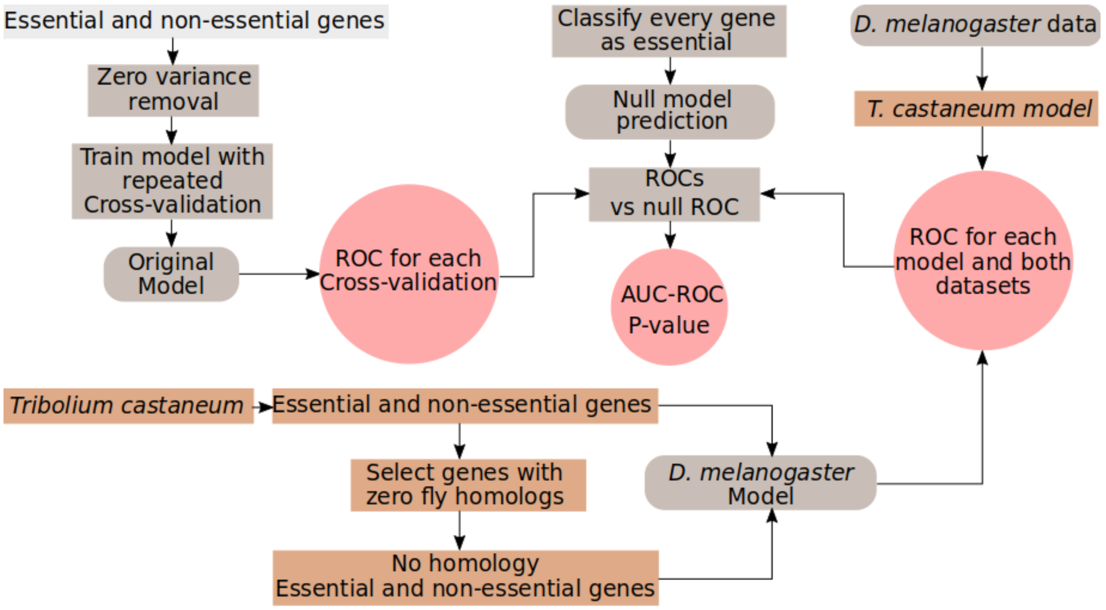

**Supplementary Figure 2**

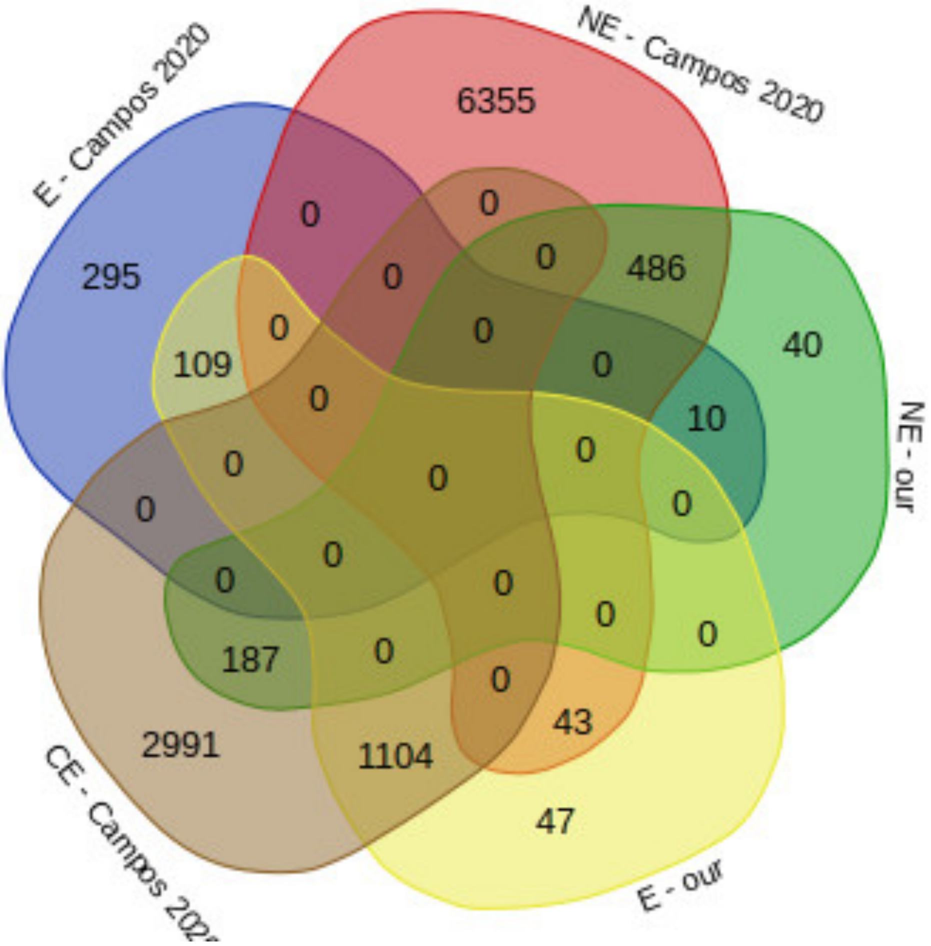

**Supplementary Table 1**

| Most frequent terms | Related records | Frequency |
|---|---|---|
| **[ empty field - no data available ]** | 1267 | 0.1274 |
| **viable** | 949 | 0.0954 |
| **recessive** | 787 | 0.0791 |
| **visible** | 698 | 0.0702 |
| **adult stage** | 690 | 0.0694 |
| **neuroanatomy defective** | 547 | 0.055 |
| **fertile** | 292 | 0.0294 |
| **third instar larval stage** | 269 | 0.027 |
| **neurophysiology defective** | 256 | 0.0257 |
| **somatic clone** | 243 | 0.0244 |

**Supplementary Table 2**

| Flybase Gene ID | Essential (0=False, 1= True) | Reference |
|---|---|---|
| **FBgn0042178** | 0 | [9] |
| **FBgn0004475** | 0 | [9] |
| **FBgn0264707** | 0 | [10] |
| **FBgn0004513** | 0 | [11] |
| **FBgn0004512** | 0 | [11] |
| **FBgn0010241** | 0 | [11] |
| **FBgn0262782** | 0 | [12] |
| **FBgn0264953** | 0 | [13] |
| **FBgn0035154** | 0 | [14] |
| **FBgn0263934** | 0 | [15] |
| **FBgn0260747** | 0 | [16] |
| **FBgn0030706** | 0 | [17] |
| **FBgn0035610** | 0 | [18] |
| **FBgn0034590** | 0 | [19] |
| **FBgn0024992** | 0 | [20] |
| **FBgn0011676** | 0 | [21] |
| **FBgn0036919** | 0 | [22] |
| **FBgn0261787** | 0 | [23] |
| **FBgn0038303** | 0 | [23] |
| **FBgn0032879** | 0 | [24] |
| **FBgn0052732** | 0 | [25] |
| **FBgn0040475** | 0 | [26] |
| **FBgn0037780** | 0 | [27] |
| **FBgn0040752** | 0 | [28] |
| **FBgn0030313** | 0 | [29] |
| **FBgn0034617** | 0 | [30] |
| **FBgn0036411** | 0 | [31] |
| **FBgn0033051** | 0 | [32] |
| **FBgn0040070** | 0 | [33] |
| **FBgn0004108** | 0 | [34] |
| **FBgn0028717** | 0 | [35] |
| **FBgn0033483** | 0 | [36] |
| **FBgn0035207** | 0 | [37] |
| **FBgn0035111** | 0 | [38] |
| **FBgn0037470** | 0 | [38] |
| **FBgn0030600** | 0 | [39] |
| **FBgn0051217** | 0 | [40] |
| **FBgn0039055** | 0 | [41] |
| **FBgn0053052** | 0 | [42] |
| **FBgn0034739** | 0 | [43] |
| **FBgn0002940** | 0 | [44] |
| **FBgn0036260** | 0 | [45] |
| **FBgn0039862** | 0 | [46] |
| **FBgn0035379** | 0 | [46] |
| **FBgn0259683** | 0 | [47] |
| **FBgn0025382** | 0 | [48] |
| **FBgn0260986** | 0 | [49] |
| **FBgn0263199** | 0 | [50] |
| **FBgn0003741** | 0 | [51] |
| **FBgn0037659** | 0 | [52] |
| **FBgn0037703** | 0 | [52] |

| | | |
|---|---|---|
| **FBgn0033233** | 0 | [52] |
| **FBgn0053182** | 0 | [52] |
| **FBgn0266570** | 0 | [52] |
| **FBgn0036366** | 0 | [52] |
| **FBgn0035166** | 0 | [52] |
| **FBgn0263025** | 0 | [52] |
| **FBgn0038948** | 0 | [52] |
| **FBgn0032671** | 0 | [52] |
| **FBgn0033238** | 0 | [53] |
| **FBgn0004050** | 0 | [54] |
| **FBgn0264272** | 0 | [55] |
| **FBgn0262369** | 0 | [56] |
| **FBgn0036125** | 0 | [57] |
| **FBgn0027528** | 0 | [58] |
| **FBgn0015575** | 0 | [59] |
| **FBgn0034135** | 0 | [60] |
| **FBgn0023517** | 0 | [61] |
| **FBgn0004575** | 0 | [62] |
| **FBgn0029976** | 0 | [63] |
| **FBgn0033744** | 0 | [64] |
| **FBgn0039666** | 0 | [65] |
| **FBgn0035847** | 0 | [8] |
| **FBgn0046885** | 0 | [8] |
| **FBgn0051438** | 0 | [8] |
| **FBgn0032439** | 0 | [8] |
| **FBgn0051882** | 0 | [8] |
| **FBgn0032754** | 0 | [8] |
| **FBgn0036970** | 0 | [8] |
| **FBgn0032585** | 0 | [8] |
| **FBgn0051406** | 0 | [8] |
| **FBgn0011832** | 0 | [8] |
| **FBgn0034156** | 0 | [8] |
| **FBgn0037974** | 0 | [8] |
| **FBgn0052301** | 0 | [8] |
| **FBgn0052282** | 0 | [8] |
| **FBgn0028987** | 0 | [8] |
| **FBgn0034427** | 0 | [8] |
| **FBgn0053462** | 0 | [8] |
| **FBgn0038299** | 0 | [8] |
| **FBgn0039739** | 0 | [8] |
| **FBgn0038888** | 0 | [8] |
| **FBgn0034870** | 0 | [8] |
| **FBgn0286516** | 1 | [66] |
| **FBgn0028418** | 1 | [18] |
| **FBgn0035416** | 1 | [67] |
| **FBgn0260859** | 1 | [67] |
| **FBgn0266722** | 1 | [67] |
| **FBgn0266724** | 1 | [67] |
| **FBgn0260655** | 1 | [67] |
| **FBgn0037551** | 1 | [68] |
| **FBgn0000723** | 1 | [69] |
| **FBgn0000463** | 1 | [70] |
| **FBgn0263864** | 1 | [71] |
| **FBgn0005654** | 1 | [72] |
| **FBgn0267975** | 1 | [73] |

| | | |
|---|---|---|
| **FBgn0032407** | 1 | [74] |
| **FBgn0266418** | 1 | [75] |
| **FBgn0027053** | 1 | [76] |
| **FBgn0000063** | 1 | [49] |
| **FBgn0000449** | 1 | [77] |
| **FBgn0260855** | 1 | [78] |
| **FBgn0260749** | 1 | [52] |
| **FBgn0036003** | 1 | [52] |
| **FBgn0000575** | 1 | [79] |
| **FBgn0025641** | 1 | [80] |
| **FBgn0003870** | 1 | [81] |
| **FBgn0260635** | 1 | [82] |
| **FBgn0002781** | 1 | [83] |
| **FBgn0264307** | 1 | [84] |
| **FBgn0263396** | 1 | [85] |
| **FBgn0031359** | 1 | [86] |
| **FBgn0015795** | 1 | [87] |
| **FBgn0020309** | 1 | [8] |
| **FBgn0020305** | 1 | [8] |
| **FBgn0021796** | 1 | [8] |
| **FBgn0004374** | 1 | [8] |
| **FBgn0267828** | 1 | [8] |
| **FBgn0031604** | 1 | [8] |
| **FBgn0003145** | 1 | [8] |
| **FBgn0004914** | 1 | [8] |
| **FBgn0000299** | 1 | [8] |
| **FBgn0001981** | 1 | [8] |
| **FBgn0032683** | 1 | [8] |
| **FBgn0261983** | 1 | [8] |
| **FBgn0040232** | 1 | [8] |
| **FBgn0023388** | 1 | [8] |
| **FBgn0000422** | 1 | [8] |
| **FBgn0263933** | 1 | [8] |
| **FBgn0040228** | 1 | [8] |
| **FBgn0014127** | 1 | [8] |
| **FBgn0086444** | 1 | [8] |
| **FBgn0002524** | 1 | [8] |
| **FBgn0000546** | 1 | [88] |
| **FBgn0003964** | 1 | [89] |

372

**Supplementary Table 3**

| Features | Overall importance | DNA/Protein feature |
|---|---|---|
| **B.DNA.twist.Roll.lag.1** | 100.00 | nuc |
| **Hartman_trans_free_energy.Protein.induced.deformability.lag.1** | 83.77 | nuc |
| **DNA.denaturation.Protein.induced.deformability.lag.1** | 83.38 | nuc |
| **prop1.Tr2332** | 80.86 | prot |
| **Stacking_energy.Tilt.lag.1** | 80.33 | nuc |
| **Twist.Propeller.twist.lag.1** | 79.56 | nuc |
| **Lisser_BZ_transition.Twist.lag.1** | 79.08 | nuc |
| **Protein.DNA.twist.Propeller.twist.lag.1** | 78.68 | nuc |
| **Aida_BA_transition.Protein.induced.deformability.lag.1** | 77.71 | nuc |
| **Twist.Protein.induced.deformability.lag.1** | 75.22 | nuc |
| **Protein.DNA.twist.Dinucleotide.GC.Content.lag.1** | 74.68 | nuc |
| **DNA.denaturation.Twist.lag.1** | 74.66 | nuc |
| **MW.Daltons.lag103** | 72.70 | nuc |
| **B.DNA.twist.Hartman_trans_free_energy.lag.1** | 71.68 | nuc |
| **ata** | 71.50 | nuc |
| **DNA.denaturation.Shift.lag.1** | 71.07 | nuc |
| **Protein.DNA.twist.Polar_interaction.lag.1** | 70.82 | nuc |
| **gt** | 70.02 | nuc |
| **cca** | 69.44 | nuc |
| **Lisser_BZ_transition.Protein.induced.deformability.lag.1** | 69.04 | nuc |

**Supplementary Table 4**

| Features | Overall importance | DNA/Protein feature |
|---|---|---|
| secondarystruct.Group2 | 100.00 | prot |
| secondarystruct.Group1 | 97.29 | prot |
| DNA.denaturation.Sugimoto_dS.lag.1 | 73.24 | nuc |
| Protein.DNA.twist.Sugimoto_dH.lag.1 | 61.74 | nuc |
| Ivanov_BA_transition.Sugimoto_dS.lag.1 | 60.95 | nuc |
| Aromatic | 56.44 | prot |
| Protein.DNA.twist.Sugimoto_dS.lag.1 | 54.96 | nuc |
| Breslauer_dS.Electron_interaction.lag.1 | 53.49 | nuc |
| Ivanov_BA_transition.Sugimoto_dH.lag.1 | 52.93 | nuc |
| prop1.Tr2332 | 50.96 | prot |
| Breslauer_dS.Propeller.twist.lag.1 | 49.90 | nuc |
| Aida_BA_transition.Sugimoto_dS.lag.1 | 49.53 | nuc |
| Breslauer_dH.Propeller.twist.lag.1 | 48.95 | nuc |
| Aida_BA_transition.Sugimoto_dH.lag.1 | 47.80 | nuc |
| Breslauer_dS.Sugimoto_dG.lag.1 | 47.02 | nuc |
| prop6.Tr2332 | 46.78 | prot |
| Bending.stiffness.Sugimoto_dS.lag.1 | 44.65 | nuc |
| DNA.denaturation.Sugimoto_dH.lag.1 | 43.37 | nuc |
| gtg | 42.90 | nuc |
| Breslauer_dH.Watson.Crick_interaction.lag.1 | 42.83 | nuc |
